# Spatial Immunophenotyping from Whole-Slide Multiplexed Tissue Imaging Using Convolutional Neural Networks

**DOI:** 10.1101/2024.08.16.608247

**Authors:** Mohammad Yosofvand, Sharon N. Edmiston, James W. Smithy, Xiyu Peng, Caroline E. Kostrzewa, Bridget Lin, Fiona Ehrich, Allison Reiner, Jayson Miedema, Andrea P. Moy, Irene Orlow, Michael A. Postow, Katherine Panageas, Venkatraman E. Seshan, Margaret K. Callahan, Nancy E. Thomas, Ronglai Shen

**Affiliations:** Department of Epidemiology and Biostatistics, Memorial Sloan Kettering Cancer Center, New York, NY; Lineberger Comprehensive Cancer Center, University of North Carolina at Chapel Hill, Chapel Hill, NC; Department of Dermatology, School of Medicine, University of North Carolina, Chapel Hill, NC; Department of Medicine, Memorial Sloan Kettering Cancer Center, New York, NY; Department of Pathology and Laboratory Medicine, Memorial Sloan Kettering Cancer Center, New York, NY; University of Connecticut School of Medicine, Farmington, CT

**Author notes:** Co-senior authors.

**Keywords:** Multiplex immunofluorescence, Deep learning, Image analysis, Spatial analysis, Melanoma

## Abstract

The multiplexed immunofluorescence (mIF) platform enables biomarker discovery through the simultaneous detection of multiple markers on a single tissue slide, offering detailed insights into intratumor heterogeneity and the tumor-immune microenvironment at spatially resolved single cell resolution. However, current mIF image analyses are labor-intensive, requiring specialized pathology expertise which limits their scalability and clinical application. To address this challenge, we developed CellGate, a deep-learning (DL) computational pipeline that provides streamlined, end-to-end whole-slide mIF image analysis including nuclei detection, cell segmentation, cell classification, and combined immuno-phenotyping across stacked images. The model was trained on over 750,000 single cell images from 34 melanomas in a retrospective cohort of patients using whole tissue sections stained for CD3, CD8, CD68, CK-SOX10, PD-1, PD-L1, and FOXP3 with manual gating and extensive pathology review. When tested on new whole mIF slides, the model demonstrated high precision-recall AUC. Further validation on whole-slide mIF images of 9 primary melanomas from an independent cohort confirmed that CellGate can reproduce expert pathology analysis with high accuracy. We show that spatial immuno-phenotyping results using CellGate provide deep insights into the immune cell topography and differences in T cell functional states and interactions with tumor cells in patients with distinct histopathology and clinical characteristics. This pipeline offers a fully automated and parallelizable computing process with substantially improved consistency for cell type classification across images, potentially enabling high throughput whole-slide mIF tissue image analysis for large-scale clinical and research applications.

## Introduction

Recent developments in multiplex spatial imaging technologies, including imaging mass cytometry (IMC), cyclic immunofluorescence (CyCIF), COdetection by inDEXing (CODEX, now PhenoCycler), multiplexed ion beam imaging (MIBI), multiplex immunofluorescence (mIF), and multiplex immunohistochemistry (mIHC), offer powerful platforms for characterization of tumor microenvironment and biomarker discovery^1–8^. These technologies enable the simultaneous detection of multiple markers of interest on a single tissue slide to gain a deeper understanding of the tumor microenvironment (TME). Among the different tissue imaging assays, mIF/mIHC carries particular significance for both research and clinical purposes. A meta-analysis by Lu et al. (2019)^9^ that included over 10 solid tumor types in 8135 patients, demonstrated that mIF/mIHC was associated with improved performance in predicting response to PD-L1/ PD-1 treatment in different solid tumor types compared to PD-L1 IHC, tumor mutational burden (TMB), or gene expression profiling [GEP, including an interferon (IFN) gamma gene signature] alone.

Most high-plex imaging studies of patient samples involve tissue microarrays (TMAs) or pre-selected regions of interest (ROIs), whereas whole-slide imaging is required to properly study intra-tumor heterogeneity and fully capture the complexity of the TME and tumor-immune cell interactions. Current whole-slide immunofluorescence imaging platforms used in clinical research include the Akoya PhenoImager HT (used for example in the AstroPath platform^3^), Ultivue, and the higher-plex platform Orion^10^. The analyses of such images can be labor-intensive, requiring specialized pathology expertise which may limit scalability in large clinical and research applications. The standard analysis involves segmenting the cells in an mIF image, examining each channel to classify marker-specific positive cells, and manually setting a marker intensity threshold (a “gating” process) for cell classification (e.g., CD3^+^ cells) on each slide. Artificial intelligence (AI) algorithms, including deep learning methods, have contributed to the task of cell segmentation in medical image analyses^11–15^. Furthermore, deep learning solutions have been proposed to perform the task of cell annotation in medical images including IF and IHC images^16–20^. However, existing methods often lack the desired precision and robustness needed for whole-slide analysis due to the varying signal intensities and background noise across the slide resulting from tissue heterogeneity, staining, and biology. Batch effects further pose challenges to achieving consistent analysis across different tissue slides, requiring all the cell annotations and biomolecular panels to be the same as the training data which limits its applications to new datasets^21^. Moreover, manual thresholds are typically set based on staining intensity averaged over a cell, losing critical information on the cellular distribution and pattern of staining (e.g., nucleus, cytoplasm, membrane). Artifacts such as segmentation errors, autofluorescence, and signal spillover can further result in incorrect cell type classifications that need manual correction. All of these analytical challenges have limited the accuracy and throughput of whole-slide tissue image analysis for large-scale applications.

To address these challenges, we developed a computational pipeline called CellGate for cell type classification and spatial immunophenotyping in whole-slide mIF images using convolution neural networks (CNNs). CNN models have been instrumental in image classification^22^, yet most applications have focused on whole-slide hematoxylin-eosin (H&E) stained tissues, including the analysis of nodal tissues to detect metastasis in breast cancer^23^ or predict mutation status from histopathology images in lung cancer^24^. Whole-slide mIF imaging is more powerful than H&E for the study of the tumor microenvironment. Here we leverage CNN architectures for automated spatial immunophenotyping from whole-slide mIF images. CellGate incorporates several major innovations. First, it is a single cell level classification by training CNN models on hundreds of thousands of labeled single-cell images to learn the variation in marker staining patterns in association with where the marker is expressed and localized and different levels of noise and background inherent to biology and chemistry. By incorporating this critical information, CellGate improves the accuracy of cell phenotyping, enabling the model to discern true signals from background and artifacts with high precision. Second, our pipeline provides a streamlined, end-to-end tool that facilitates high-quality fully automated image analysis eliminating the bottleneck of human annotation. We demonstrate the applicability of CellGate to two mIF imaging datasets from different patient cohorts, independently performed at different institutions.

## Results

### CellGate, a deep learning pipeline for automated and high-precision spatial immuno-phenotyping from mIF images

CellGate is a deep learning computational pipeline to automate whole-slide mIF tissue imaging analysis offering high-quality and streamlined cell classification and spatial immuno-phenotyping. Figure 1 illustrates the workflow of CellGate. The process begins with a patching algorithm, where the pipeline divides the large whole-slide mIF image (tens of thousands of pixels in both height and width) into smaller patches for more efficient processing. Next, the CellGate pipeline employs the StarDist model^14^ to segment the cell nuclei on the DAPI-stained channel for each patch. Since DAPI is a clear and crisply stained nuclei marker, using it to identify and segment the cell nuclei eliminates the parts of the slides with artifacts and other unwanted background, simplifying the process. The spatial location and distribution of the segmented cell nuclei are preserved during this step. Using the segmentation mask, cells are then cropped out into single-cell images, resized to 64×64 pixels dimension, and fed into pre-trained CNN models. This model generates classification labels (positive or negative), and classification probabilities for each of the 8 marker-stained channels: panCK-Sox10, CD3, CD4, CD8, CD68, FOXP3, PD-1, and PD-L1. An illustration of positive marker staining in an area with the presence of positively stained cells is shown in Figure 2A, with Figure 2B showing the positive staining at a single cell level in the original grayscale image. The predicted labels and the probabilities of the positive prediction are generated for each cell across all patches to complete a whole-slide single cell classification. In the subsequent step, the CellGate pipeline combines the label prediction for these markers to predict a cell phenotype (e.g., exhausted T-cell) (Figure 1). The spatial locations and the predicted phenotypes of the cells are then used to create a pseudo-plot of the cancer tissue, allowing for the study of the spatial distribution of different cells within the tumor microenvironment.

**Figure 1.**
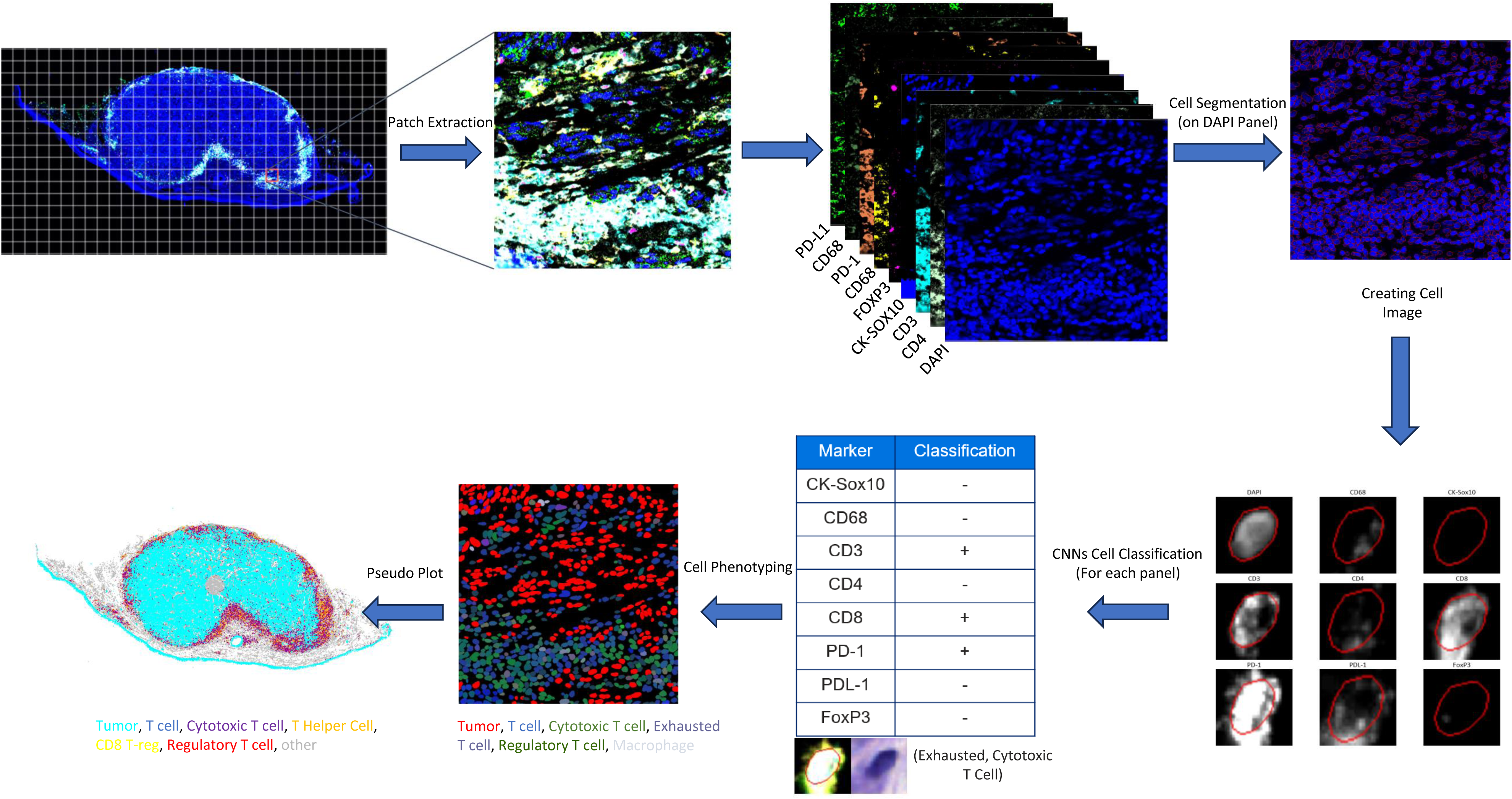
CellGate pipeline. The process starts with dividing the whole-slide image into patches, and cell segmentation based on the DAPI channel, generating single cell images from the segmentation mask, followed by a series of gating and phenotyping steps. Specifically, the single cell images are fed into pre-trained convolutional neural network models to predict class label prediction for each marker (e.g., CD3+ vs CD3-). Then the predictions are stacked across all the marker channels to obtain combined phenotype (e.g., CD3+CD8+PD-1+, exhausted cytotoxic T cell). The location coordinates can then be used to map the phenotyped cells onto a pseudo-plot to study the tumor immune microenvironment.

**Figure 2.**
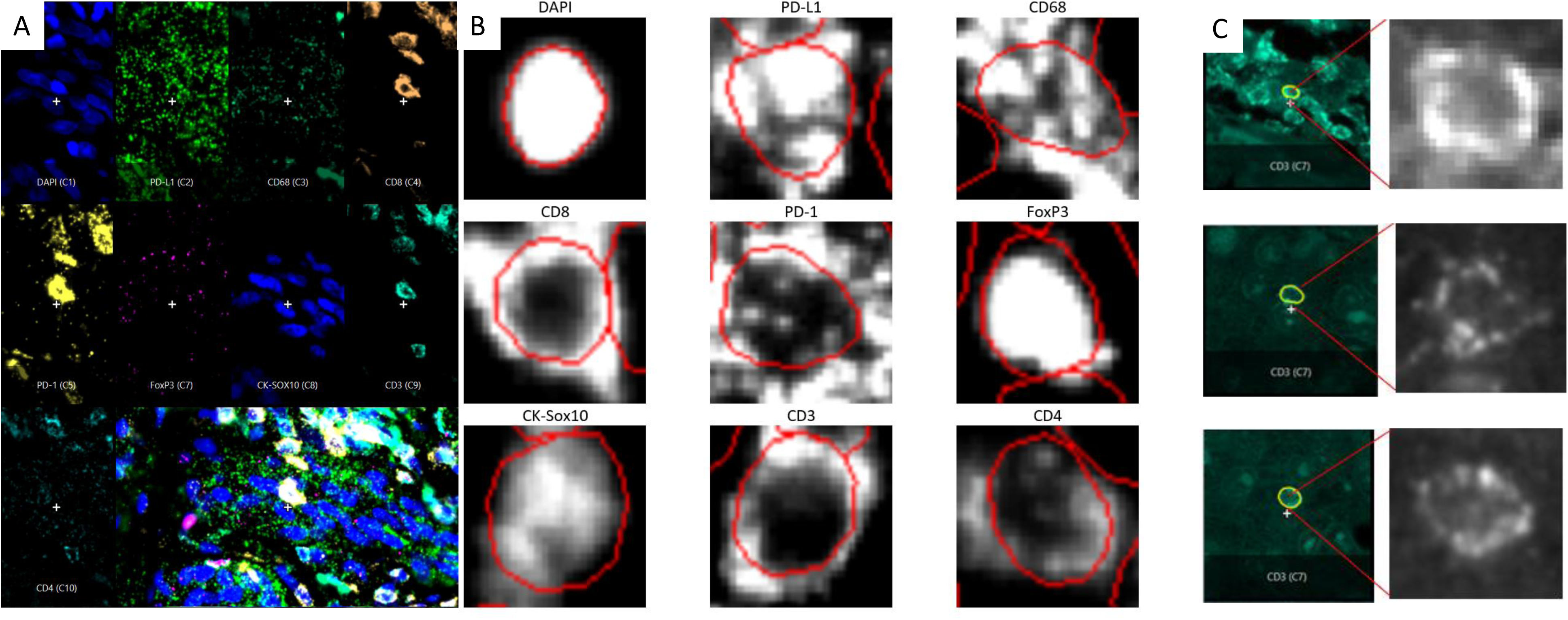
A typical view of an mIF slide with different panels is displayed separately. A. Illustration of positive marker staining in a representative area of mIF images with the presence of positively stained cells in each marker channel. B. Marker staining pattern at the single cell level. C. An example of a weakly stained CD3 with dim intensity but strong cellular pattern picked up by CellGate.

### Training the CNN model

CellGate uses a combination of CNNs with a modified VGG19 network^25^ as the backbone model. Each CNN model is independently trained on each marker. The input image for the VGG19 network was resized to 64×64×3 and pre-trained weights from the ImageNet dataset^26^ in the Keras library in TensorFlow^27^ were used to start the training process. The 16-bit grayscale cell images from the marker panels were expanded to a 3-layer image to match the model input before being fed into the VGG19 network. The output layer of the VGG19 model was then flattened and a dense layer with 2 units and softmax activation was used as binary classification for each panel. More details can be found in the Methods section.

To train the model, we used a dataset consisting of non-lymph node metastatic samples from 34 patients with unresectable melanoma treated at Memorial Sloan Kettering Cancer Center (MSK) with archival pre-treatment tumor specimens available^28^. Whole tissue sections were stained using Ultivue UltiMapper I/O Immuno8 Kit (Cambridge, MA, USA) containing CD8, PD-1, PD-L1, CD68, CD3, CD20, FOXP3, and pancytokeratin □+ □SOX10 as described in Vanguri et al. (2023)^29^. Whole slide imaging was acquired with a Zeiss AXIO Scanner. The image analysis of the 34 MSK samples was manually performed by a dedicated pathology technician, with manual annotation and extensive quality control by the clinical and pathology team. The final labeled data included the coordinates for all cells from each mIF image and the positive/negative classification for each cell for each marker.

Using this manually curated dataset as labeled data, we trained CNN models for automated gating and immunophenotyping at the single cell level. We focused on 34 non-lymph node samples from MSK, dividing them into a training set (N=29), and a test set (N=5). For the training set, we used cell location and bounding box coordinates to crop the manually labeled single-cell images for model training. For each marker, a fraction of the positive cells were randomly selected such that the training image dataset contained at least 50,000 positive cells. An equal number of cells with negative labels from the same mIF slide were randomly selected to form the negative controls. This sampling method is necessary to address cell type imbalance during model training. The CellGate pipeline was trained on over 750,000 single cells in total across the 29 stacked images. A binary cross entropy loss function was employed to optimize the CNN models. The CNN models were trained for 200 epochs with no early stopping, and different learning rates and optimizer functions were implemented to fine-tune the model for each marker. To further generalize the trained models, cross-validation training was done on different sets of training and test mIF images in which the model maintained its performance across different markers, indicating the accurate and consistent performance of the developed pipeline. Table 1 shows the training set cross-validation accuracies, averaged over 5 different training and testing combinations of the 34 mIF whole slides.

**Table 1.** Average cross-validation accuracies for different sample selections.

| Biomarker Panel | CD3 | CD8 | CD68 | CK-Sox10 | FOXP3 | PD-1 | PD-L1 |
| --- | --- | --- | --- | --- | --- | --- | --- |
| Average Accuracy | 83.3 | 88.7 | 76.5 | 78.1 | 89.4 | 66.3 | 77.1 |

A total of 114,108 positive and negative cells (57054 each) were used to train the model to classify CD3 positive cells, achieving a test accuracy of 83.3% (Table 1). The CD8 marker model was trained on a total of 98,934 positive and negative cells (49197 each) and achieved a higher test accuracy of 88.7%. The training accuracy varies based on the pixel intensity distributions of the marker. Generally, the CD3 stain has more noise across the slide and within the cell boundaries compared to CD8, which shows a crisper staining pattern with the positive staining signals distributed closer around the nuclei. The CK-SOX10 marker is a dual channel, with the cytokeratin (CK) staining epithelial cells and SOX10, a lineage marker for melanoma, defining tumor cells. The CNN model for CK-SOX10 was trained on 110,470 positive and negative cells (55235 each), achieving a training accuracy for the CK-SOX10 of 78.1%.

The CD68 channel identifies macrophages in the tumor microenvironment and exhibits more artifactual staining, noise, and spillover stains (Figure 2A) compared to CD3 and CD8. We trained the CNN model for the CD68 biomarker with 108,436 positive and negative cells (54218 each). The model learned the intensity distribution for CD68 positive cells and achieved a test accuracy of 76.5%. The CNN models for PD-1 and PD-L1 achieved 66.3% and 77.1% accuracy, respectively. Both PD-1 and PD-L1 stains show a more varied pattern of staining (Figure 2B) and higher levels of scattered noise across the slide, making classification more challenging. FOXP3+ cells were the least expressed positive cells in our training dataset, with only about 136,000 cells labeled across the 29 mIF slides. In total 108,622 FOXP3 positive and negative cells (54,311 each) were used to train and test the CNN model, resulting in a training accuracy of 89.4%.

### CellGate can reproduce cell classification labels from expert pathology with a high precision-recall

We then assessed the model performance on whole-slide analyses of the 5 unseen test samples, set aside initially. These slides are called “unseen test slides” as no training was performed on them, allowing for an unbiased assessment of cell class prediction accuracy. To test the robustness of the algorithm, we chose the test tissue samples with a varying total number of cells, ranging from under 10k to over 400k, and a wide range in the proportion of immune cells (e.g., from ∼1% to ∼30% CD3+ cells, Supplementary Table 1A).

When evaluating performance on a whole slide where the cell type proportions are highly imbalanced, precision-recall is an appropriate metric of evaluation. When applied to the 5 unseen mIF whole slides, the model achieved an average 0.90 precision-recall area under the curve (PR-AUC) ranging between 0.81 and 0.95 for CD3 (Table 2, Supplementary Figure 1A). Sensitivity, specificity, F-score, positive predictive value (PPV), and negative predictive value (NPV) using a 0.5 probability cutoff for binary prediction presented in Supplementary Table 1A, along with the total number of cells for each slide, the percentage of CD3+ cells by pathologists’ gating, and the predicted proportion of the CD3+ cells. The R-squared value for the corresponding predicted CD3+ cells and the CD3+ percentages gated by pathologists was 0.98 (Figure 3). The PR-AUC for CD8 was 0.86 (range: 0.77-0.94). Since CD8+ cells are a subset of CD3+ cells, slightly lower PR-AUC is expected for subsets with lower frequency.

**Figure 3.**
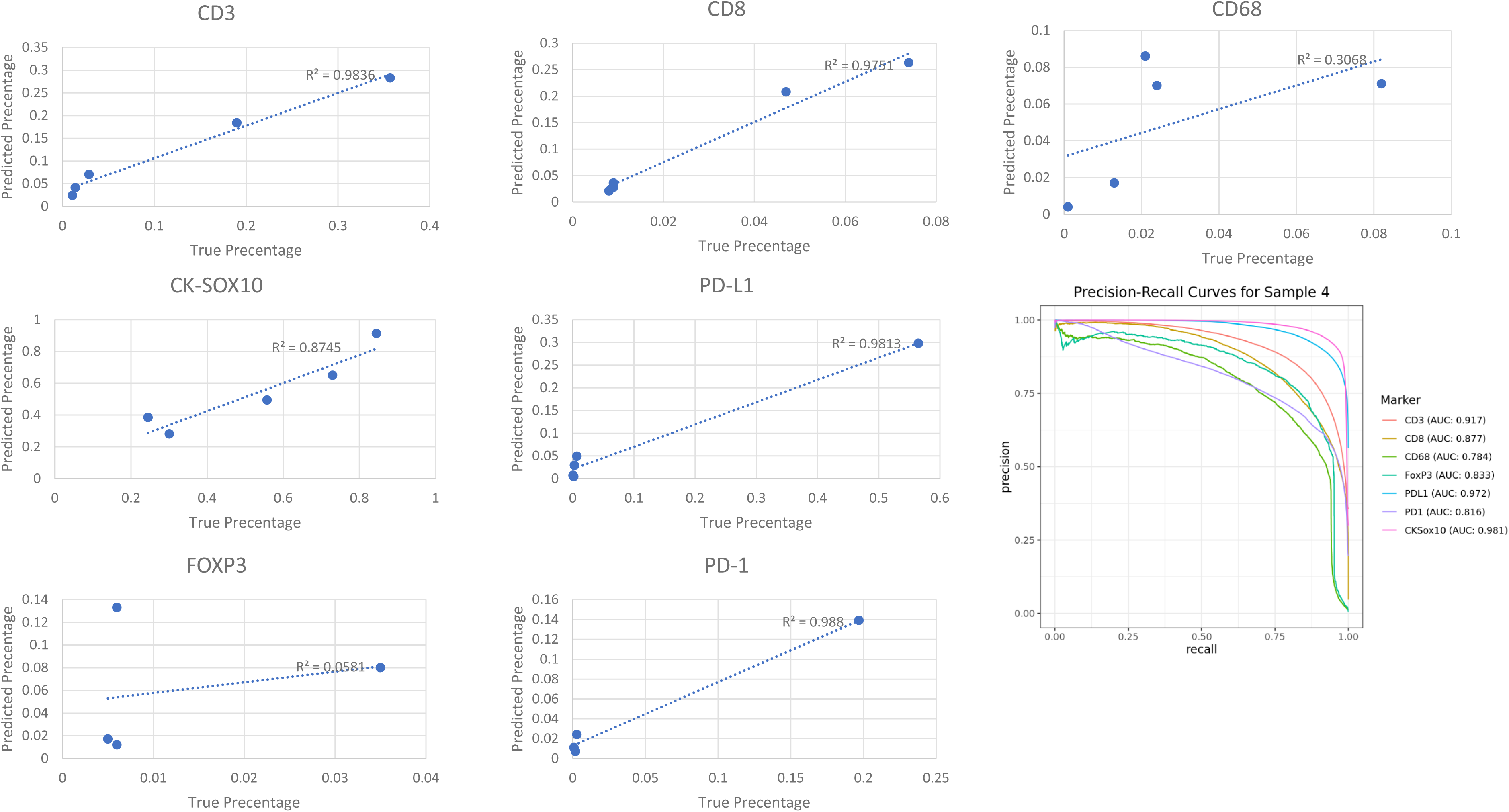
Fraction of positive cells for predicted labels compared to the manual (true) labels and the precision-recall curves for different biomarkers for the testing mIF whole slide sample 4.

**Table 2.**
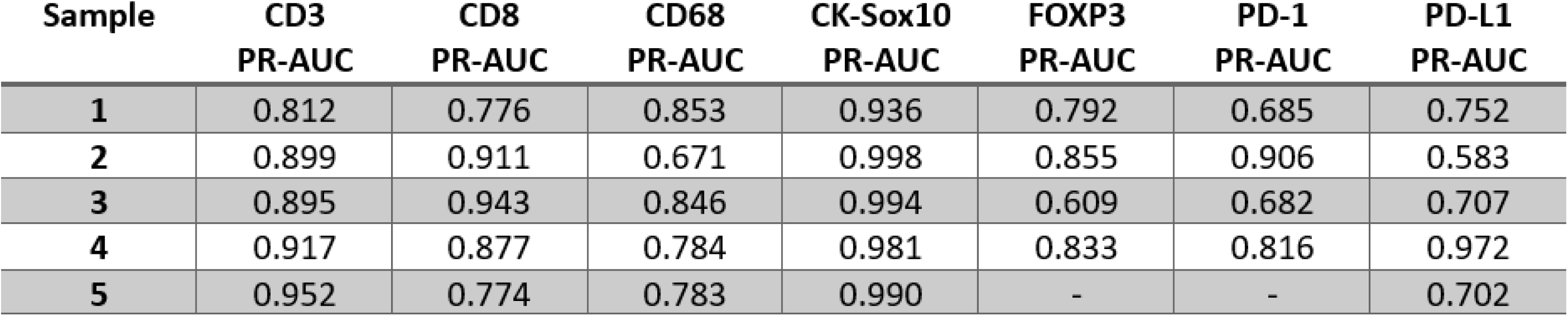
Precision-Recall AUC for each biomarker for the testing mIF whole slide images.

| Sample | CD3<br>PR-AUC | CD8<br>PR-AUC | CD68<br>PR-AUC | CK-Sox10<br>PR-AUC | FOXP3<br>PR-AUC | PD-1<br>PR-AUC | PD-L1<br>PR-AUC |
| --- | --- | --- | --- | --- | --- | --- | --- |
| 1 | 0.812 | 0.776 | 0.853 | 0.936 | 0.792 | 0.685 | 0.752 |
| 2 | 0.899 | 0.911 | 0.671 | 0.998 | 0.855 | 0.906 | 0.583 |
| 3 | 0.895 | 0.943 | 0.846 | 0.994 | 0.609 | 0.682 | 0.707 |
| 4 | 0.917 | 0.877 | 0.784 | 0.981 | 0.833 | 0.816 | 0.972 |
| 5 | 0.952 | 0.774 | 0.783 | 0.990 | - | - | 0.702 |

Furthermore, the CellGate pipeline can potentially improve the standard manual gating approach (based on intensity thresholds) by further incorporating the spatial pattern of the marker staining on cells. The manual intensity thresholding may misclassify some cells with dimmer pixels as negative for that marker, whereas CellGate can identify these as T cells by recognizing the circular staining pattern of CD3 that outlines the cell (Figure 2C). We illustrate such cases where the cell was initially classified as negative for CD3 cells using the manual gating threshold and were reclassified as positive CD3 cells by CellGate, potentially increasing the gating sensitivity (Figure 2C).

The results for CK-SOX10 from the CNN on the 5 test/new slides had the highest precision-recall AUCs among all channels, with an average PR-AUC of 0.98 (range:0.94-0.99). The tumor nuclei on the CK-SOX10 channel display a crisp nucleus staining and a uniform pixel distribution inside the positive cells (Figure 2B). For CD68, the performance on the unseen whole slides achieved an average PR-AUC of 0.79, ranging between 0.67, and 0.85, with the lowest PR-AUC from a sample with less than 1% of the cells classified as CD68+ cells by the pathologists’ annotations. Even though the percentage of predicted positive cells by the CNN model is very close to the one from the gated dataset (both less than 0.5%), it is expected to have a lower PR-AUC value for rare biomarkers. This is true for FOXP3, PD-1, and PD-L1 as well. The PR-AUC values for each of the 5 test slides are presented in Table 2 and the rest of the evaluation metrics are listed in Supplementary Table 1. We also evaluated overall concordance in the estimated percentages of the predicted positive cells from the model compared to those labeled by pathologists (true labels) at the whole-slide level as shown in Figure 3. The concordance as measured by R-squared is very high for CD8, CD3, PD-1, PD-L1, and CK-SOX10. The concordance is reduced for CD68 likely due to the more challenging staining pattern. The concordance for FOXP3 is low because the frequency of FOXP3+ cells on these 5 test slides is very low (less than 1% in most cases).

Overall, the results demonstrated that our fully automated and parallelizable computational pipeline can reproduce expert pathology analysis with high accuracy. This capability potentially enables scalable mIF tissue whole slide image (WSI) analysis, making it a powerful tool for large-scale clinical and research applications.

### Whole-slide spatial immunophenotyping using CellGate in an independent cohort of primary melanoma patient samples

Next, to further assess the applicability of CellGate on mIF imaging data independently generated at a different institution, we applied the pre-trained CellGate pipeline on mIF whole-slide tissue images of 9 primary melanoma samples from the UNC pathology archive. Using the Ultivue Immuno8 FixVUE assay, eight immune-related and tumor markers (CD3, CD4, CD8, CD68, FOXP3, PD-1, PD-L1, CK-SOX10) were stained at UNC to identify and characterize T-cells subsets, immunosuppressive and immune checkpoint markers in the melanoma TME. Given raw single cell images cropped from the corresponding cell segmentation mask, the CellGate image model returns the predicted cell classification for each marker. The stacked prediction then leads to the phenotypic classification of the corresponding cell, focusing on 18 major cell types including tumor cells (SOX10+), immune-evading tumor cells (SOX10+PD-L1+), FOXP3+ tumor cells, CD68+ tumor cells, immunosuppressive macrophages (CD68+PD-L1+), macrophages (CD68+), CD4/CD8 double positive T cells (CD3+CD4+CD8+), CD4/CD8 double positive FOXP3+ T cells, exhausted CD4/CD8 double positive T cells (CD3+CD4+CD8+PD-1+), exhausted T helper cells (CD3+CD4+PD-1+), exhausted T cells (CD3+PD-1+), exhausted cytotoxic T cells (CD3+CD8+PD-1+), CD8+ regulatory T cells (CD3+CD8+FOXP3+), regulatory T cells (CD3+CD4+FOXP3+), FOXP3+ T cells (CD3+FOXP3+), T cells (CD3+), T helper cells (CD3+CD4+), cytotoxic T cell (CD3+CD8+). Figure 4 displays the cell type composition in each of the UNC samples, with 4A showing the proportion of major cell types (tumor cells, T cells, macrophages) quantified from the whole-slide image and 4B further delineating the functional subsets within the tumor microenvironment.

**Figure 4.**
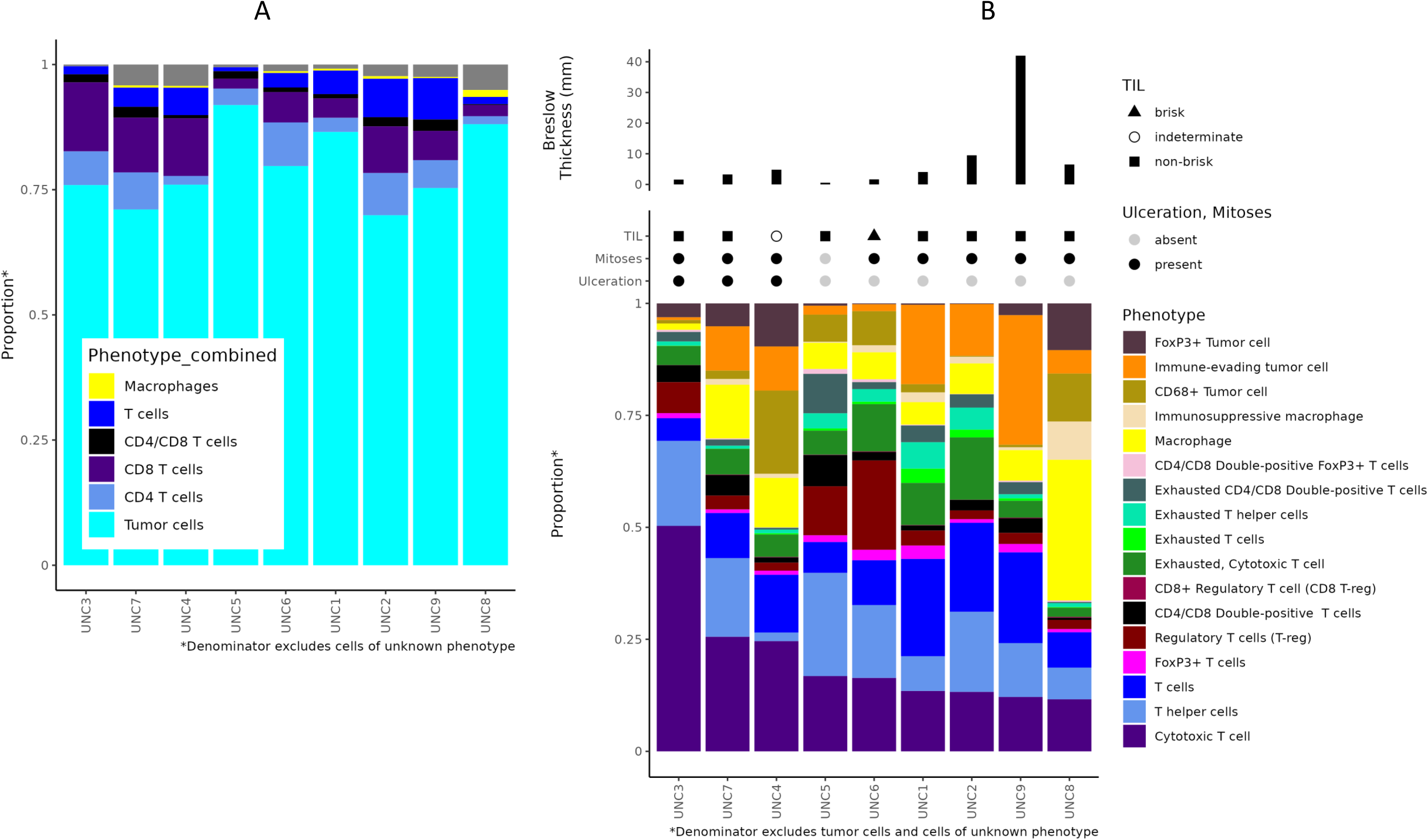
Stacked bar plots of cell type composition in the validation UNC primary melanoma samples. A. Proportion of major cell types including tumor cell, T cell, and macrophage calculated from CellGate classification of all segmented cells based on DAPI stain from each whole-slide image analysis. B. Composition of the 17 functional subsets of tumor and immune cells of interest in each tumor sample. Proportion is calculated as the total count of each cell type divided by the sum of the 17 cell types.

Tumor-infiltrating lymphocytes (TILs) have long been recognized as a biomarker of host-immune response and crucial in anti-tumor immunity and the presence of TILs around and inside the tumor carries prognostic value^30^. In primary melanoma, the histopathologic measurement of TILs, or TIL grade, is evaluated by a pathologist using melanoma tissue sections stained with H&E and scored as absent, non-brisk, or brisk. Primary melanoma TIL grade has prognostic value but major limitations include interobserver differences in scoring and limited TIL quantitative and spatial information. Also, while major immune cell types, their subpopulations, and their functional states have biological and clinical relevance^3,31–33^, these features are better assessed with cellular marker staining using whole-slide tissue imaging.

We note in Figure 4B that 7 of the 9 UNC samples had a non-brisk TIL grade, yet the spatial immuno-phenotyping from the mIF images revealed a great heterogeneity in the compositional differences in lineage and functional cell states, and potential tumor-immune cell interactions in spatial niches. Figure 5 presents mIF analysis of a validation sample of primary melanoma (UNC1), demonstrating that T cells are primarily located within 150 µm (on average) from the tumor margin. This finding is consistent with the non-brisk TIL classification (focal distribution of infiltrates) evaluated by pathologists on H&E. However, the mIF analysis revealed a much deeper understanding of the spatial distribution of T cell functional subsets. The magnified mIF image and corresponding pseudo plot of a local region reveal a dense mix of cytotoxic CD8, T helper, and Treg cells, but primarily located in the tumor margin suggesting an exclusive immune cell topography. Interestingly, a proportion of double positive T cells (CD8+CD4+, dark green dots in pseudo plot) is observed in the mix. Double positive (DP) T cells are often considered an anomaly, generally overlooked as cell doublets or cells having escaped faulty thymic selection. However, recent studies have shown its relevance in tumors and demonstrated its origination from antigen stimulated single positive (CD8+ or CD4+) T cells differentiating into DP T cell subsets gaining in polyfunctional characteristics and displaying enhanced cytotoxic potential^34^. Figure 5 also shows a summary of all 18 major cell type frequencies in the bar chart, a spatial UMAP showing the T cells (blue) are close to tumor cells (teal) yet not intermixed. Pseudo plots of binary cell classification for each of the 8 markers on the whole-slide of UNC sample 1 are provided in Supplementary Figure 2, with the combined phenotyping shown in Supplementary Figure 3.

**Figure 5.**
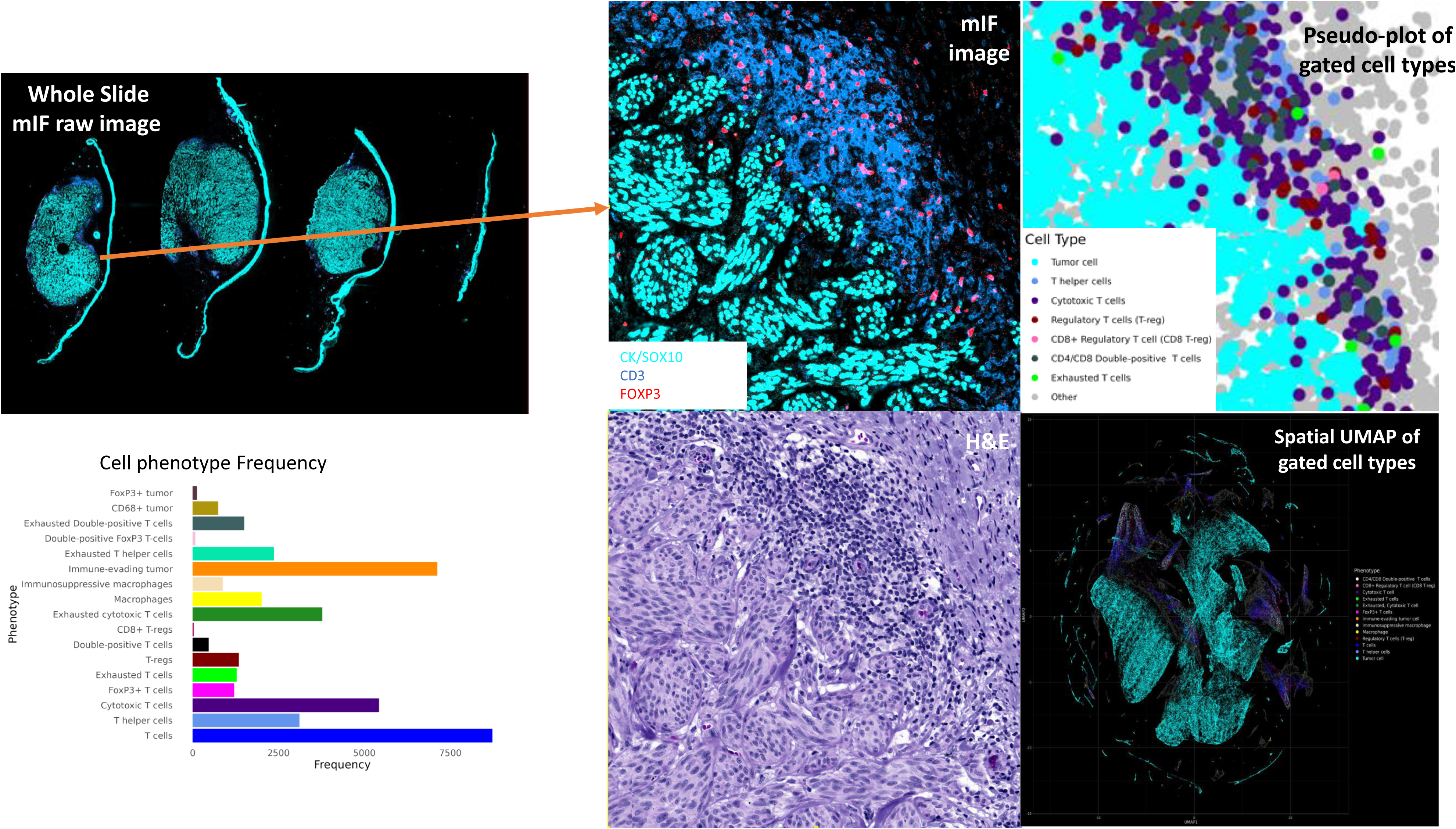
Validation sample 1. The top left panel shows the whole-slide mIF raw image with four pieces of tumor sampled from a patient with primary melanoma. Clinicopathological features of this case include non-brisk TIL, Breslow 4.05mm, and spindle cell histology. The top right panel shows an insert of a small area displaying a complex mix of different immune cell types primarily located within 150 μm of the tumor margin. The cell type frequency from whole-slide gated results is summarized in the bar-chart and the spatial UMAP at the lower right corner.

Figure 6 presents a second case also classified as non-brisk TIL grade, yet it shows a very different immune cell topography. In this case, PD-1 expressing T cells cluster around the fibrosis tracks within the tumor bulk, with nearby tumor cells highly expressing PD-L1. This striking example of tumor-immune cell interactions along the PD-1/PD-L1 axis highlights inhibitory signals that potentially modulate the balance between T-cell activation, tolerance, and immunopathology. The fibrotic pattern and cellular interactions are apparent from the raw image and the corresponding pseudo-plot of a representative local region in Figure 6. The second case has a high Breslow thickness (9.5 mm). The spatial UMAP shows a greater intermixing of the immune and tumor cells compared to UNC1.

**Figure 6.**
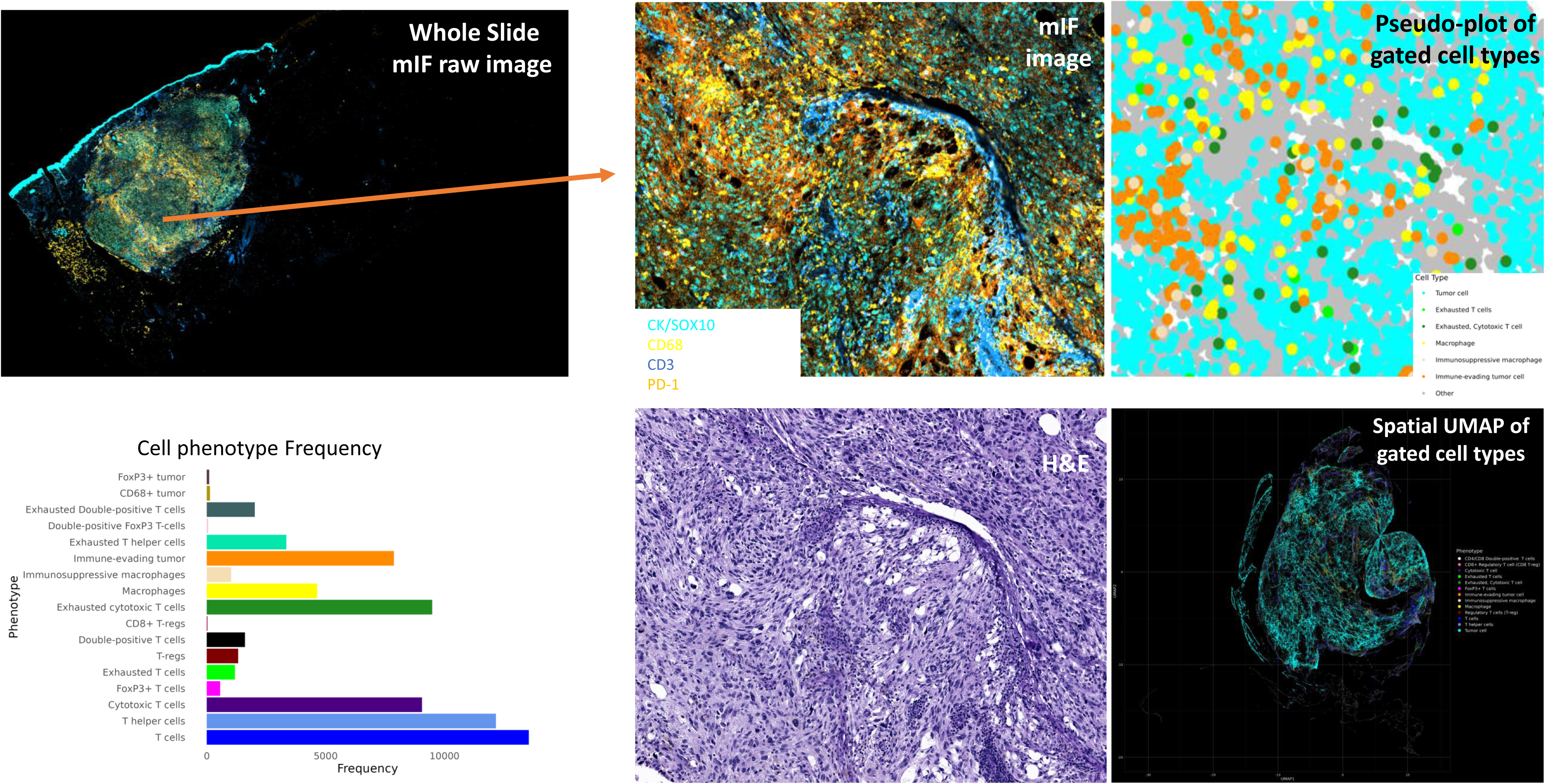
Validation sample 2. The top left panel shows the whole-slide mIF raw image of the tumor sample from a patient with primary melanoma with clinicopathological features including non-brisk TIL, Breslow 9.5, and nodular pattern. The top right panel shows an insert of a small area displaying clustering of PD-1 expression T cells along the fibrosis track and PD-L1 expressing tumor cells in close proximity. The cell type frequency from whole-slide gated results is summarized in the bar-chart and the spatial UMAP at the lower right corner suggesting a more intermixed tumor and immune cell composition.

Gated cell type frequencies and pseudo plots showing the spatial distributions of the cell types for the other validation UNC samples are shown in Supplementary Figure 3. The pseudo-plots show a concordant representation of the raw images, capturing the tumor areas quite well. The cell type frequency bar chart also allows a quick view of the differences in immune cell composition across samples. For example, the ratio of CD8 over CD4 (cytotoxic over T helper and/or Treg), the extent of T-cell exhaustion, the enrichment of immunosuppressive subsets (e.g., PD-L1 expressing macrophage) or immune-evading tumor cells (PD-L1 expressing tumor cells), along with the spatial coordinates of gated cell phenotype can greatly facilitate downstream analysis to identify spatial TME features to correlate with histopathological features and clinical outcome.

Overall, our validation analysis confirmed that CellGate can reproduce expert pathology analysis of whole-slide tissue imaging with high precision in a fully automated way. It serves as a powerful tool for in-depth analysis of cell-cell interactions in the TME.

## Discussion

In this study, we present CellGate, a deep learning computational pipeline for fully automated spatial immunophenotyping of whole-slide multiplexed tissue images. The model is currently trained on a set of mIF whole slide images of melanoma samples with the immuno8 panel labeled at the single cell level. We show the model has high accuracy for classifying melanoma cells stained with SOX10, and immune cell markers CD3 and CD8. In some cases, CellGate results show enhanced sensitivity in capturing marker positive cells by accounting for the spatial distribution of the stain. An illustrative example in Supplementary Figure 4A demonstrates that CellGate captured positive CD8 cells missed by the original gating in a zoomed-in region. This increased sensitivity in capturing true positives in multiple representative regions from the whole-slide mIF image was manually reviewed and confirmed. As a result, CellGate classified more CD8 positive cells for the 5 test slides compared to the original gating and the percentage of CD8 positive cells is higher (Figure 3). The classification concordance for CD68 is reduced likely due to a higher level of spillover, background, and artifactual staining across the slide, and the lack of specificity of this marker, as CD68 can also be found in fibroblasts, endothelial cells, and tumor cells. To improve the accuracy of classifying macrophages, the inclusion of other markers in combination with CD68 may be needed. The PD-L1 biomarker shows the most scattered staining beyond cell boundaries, making a cell-focused gating strategy challenging. In a zoomed-in view of a local region from a whole-slide image, Supplementary Figure 4B illustrates that the model detects fewer positive PD-L1 cells compared to the original gating labels (cells in red square). Using manual thresholding is subject to classifying PD-L1 positive cells that could contain background stains, and CellGate is more robust against these background signals by using a single cell image-based gating approach. Furthermore, studies have shown the detection of PD-L1 staining in tumor-derived extracellular vesicles (TEV) ^35–38^, suggesting some of the inter-cellular staining may be a true signal but a TEV marker may be needed. FOXP3 shows the most discordance between the labeled data and the prediction, likely due to the challenge of labeling infrequent cell types, as it had the least number of expressed cells in the trained marker set. An improved label-generation process involving both human curation and innovative computational algorithms is needed to improve the performance for classifying rare cell types. Our model will be further fine-tuned and adapted with additional datasets.

The key concept of CellGate is to use CNN image models to learn cellular staining patterns for cell type classification, providing robustness against variations in staining intensity and background noise within and between slides. This approach can be broadly applied beyond immunofluorescence, given a set of labeled datasets for the specific marker and platform to retrain the model. However, the performance of such models trained for analyzing other marker, chemistry, and the more recent developments in 3D molecular tissue imaging^39,40^ remains to be seen. In addition, generating human-labeled data is costly, and automated label generation may be required for extended applications.

Multiplexed tissue imaging carries the promise of new biomarker discovery in the context of immuno-oncology. Recent reports on multiplexed tissue imaging in immuno-oncology settings have led to novel discoveries of functional T cell subsets and their spatial distribution in the TME with immunological and clinical relevance. For example, Tumeh et al. (2014)^31^ analyzed samples from 46 patients with metastatic melanoma obtained before and during anti-PD-1 therapy (pembrolizumab) using mIHC. They showed pre-existing CD8+ T cells distinctly at the invasive tumor margin are associated with expression of the PD-1/PD-L1 immune inhibitory axis and may predict response to therapy. Berry et al. (2021)^3^ performed a 6-plex whole-slide mIF tissue imaging of pre-treatment FFPE tumor specimens from 53 patients with metastatic melanoma. They found high density of CD8+FOXP3+PD-1low/mid cells was closely associated with response to PD-1 blockade. Antoranz et al. (2022)^41^ found the spatial relationship between cytotoxic T cells and PD-L1 expressing macrophages was predictive of ICI response in a small cohort of melanoma patients. Attirill et al. (2022)^33^ stained FFPE primary melanoma tissue using mIHC from 66 stage II melanoma patients with available clinical and follow-up data. They reported a CD39+ tumor-resident CD8+ T-cell subset enriched for PD-1 expression comprised a significantly higher proportion of intratumoral and stromal CD8+ T-cells in patients with better recurrence-free survival. In cutaneous T cell lymphomas (CTCL), Phillips et al. (2021)^42^ found the spatial organization of PD-1+ CD4+ T cells and tumor cells predicted response to pembrolizumab. Among the most promising emerging spatial biomarkers are tertiary lymphoid structures (TLS) which are tumor-associated B cell aggregates that may function as micro-lymphoid organs playing a role in antigen presentation and T-cell priming^43–45^.

To translate these findings into clinical trial applications and personalized immunotherapy programs, larger validation studies are required. Here we report a newly developed deep learning computational pipeline for spatial immuno-phenotyping and high-quality fully automated image analysis to facilitate large-scale tissue imaging studies. Furthermore, by applying CellGate to a large number of mIF whole-slide images, we can build an atlas of phenotyped cells and apply these phenotype labels in co-registered H&E images. This, in turn, will enable training models on these cell labels using H&Es, ultimately allowing cell phenotypes to be determined directly from H&E images. Therefore, CellGate could have much broader applicability, considering the wide availability and utilization of H&Es in routine diagnosis. To translate our findings into clinical trial applications and personalized immunotherapy programs, larger validation studies are required.

## Methods

### Datasets

#### MSKCC dataset

A total of 34 patients with unresectable melanoma were treated at Memorial Sloan Kettering Cancer Center (MSKCC) and had available archival pre-treatment tumor specimens. Whole tissue sections were stained using Ultivue UltiMapper I/O Immuno8 Kit (Cambridge, MA, USA) containing CD8, PD-1, PD-L1, CD68, CD3, CD20, FOXP3, and pancytokeratin □+ □SOX10 followed by opal tyramide staining containing TCF1/7, TOX, Ki67, LAG-3 as described in Vanguri et al. (2023)^29^. Whole slide imaging was acquired with a Zeiss AXIO Scanner. The images from three rounds of staining were stacked with Ultivue Ultistacker v1.0b6 software to get the final 12-marker stacked images which can be visible and analyzable in HALO software (Indica Labs, Albuquerque, NM). Stacked TIFF images were reviewed and analyzed in HALO. Areas of necrotic or folded tissue were manually excluded, and representative stroma, tumor, and glass regions were manually selected to inform a regional classifier. The AI module in the OPAL protocol was used to perform nuclear segmentation based on the Hoechst-stained mask. Whole cells were defined by dilating the nuclear segmentation by 1.2 microns beyond the Hoechst signal, and each cell was assigned a unique ID. Individual marker channels were manually gated by a pathologist technician using the mean intensity via the HALO HighPlex FL 4.1.3 module. This gating process classifies positive cells from background staining based on a visually determined intensity threshold by a dedicated pathology technician. After several rounds of visual inspection, manual annotation, and extensive quality control by the pathology team, the final labeled data for all cells from whole slide images was generated. This cell-level dataset included the coordinates for all cells from each mIF image as well as the positive or negative classification for each cell for each marker.

#### UNC dataset

Eight primary melanoma samples were obtained from the University of North Carolina (UNC) pathology archive. Ultivue FixVUE Immuno8-profiling is a multiplex immunofluorescent assay allowing co-localization detection of multiple proteins for expression within the tumor microenvironment. Using the 8-Plex Immune-profiling panel (Ultivue Immuno8 FixVUE), we stained primary melanoma tissues for 8 immune-related and tumor markers (CD3, CD4, CD8, CD68, FOXP3, PD-1, PD-L1, CK/SOX10) to identify and characterize T-cells subsets, immunosuppressive and immune checkpoint markers in the melanoma tumor microenvironment. Individual and co-expression of the markers provides the ability to distinguish multiple cellular phenotypes [T cells (CD3+), T helper cells (CD3+CD4+), cytotoxic T cells (CD3+CD8+), exhausted T cells (CD3+PD-1+), exhausted cytotoxic T cells (CD3+CD8+PD-1+), T-regs (CD3+CD4+FOXP3+), FOXP3 T cells (CD3+FOXP3+), macrophages (CD68+), immunosuppressive macrophages (CD68+PD-L1+), double-positive T cells (CD3+CD8+CD4+), CD8+ regulatory T cells (CD3+CD8+FOXP3+), tumor cells (CK-SOX10+), and immune-evading tumor cells (CK-SOX10+PD-L1+)]. Formalin-fixed, paraffin-embedded primary melanoma tissues are sectioned (5 microns) and stained with the 8-Plex Immune-profiling panel (Ultivue Immuno8 FixVUE). Staining was performed in the Leica Bond Rx fully automated research stainer (Leica Biosystems) according to the Ultivue protocol. Briefly, slides were dewaxed in Bond dewax solution (AR9222) and hydrated in Bond wash solution (AR9590). Heat induced antigen retrieval was performed for 20 min at 100°C in Bond-Epitope Retrieval solution 2 pH-9.0 (AR9640). All the following steps were done at the ambient temperature. After pretreatment, the oligo-labeled UltiMapper® Antibody Immuno8 mix was added as a single step for 1h followed by the UltiMapper pre-amplification (25 min), in situ-Plex PCR amplification (90 min) and counter stained with UltiMapper® Nuclear Counter stain for 15 minutes. UltiMapper® fluorescent probes (FITC, Cy3, Cy5, and Cy7) were incubated for 25 minutes for visualization. Four antibodies and the nuclear counter stain were imaged at 20x magnification in 5 channels (DAPI, FITC, Cy3, Cy5, and Cy7) using a ZEISS AxioScan Z1 whole slide scanner followed by the UltiMapper® Exchange protocol to remove the first fluorescent probes and hybridize the second fluorescent probe set. The second set of four fluorescent probes was imaged and then the two images were co-registered with the Ultivue Ultistacker software. DAPI co-registration was used as a quality measure of alignment. Washes will be done using Bond Wash (AR9590) and the UltiMapper® wash solutions. After all markers are visualized and scanned, fluorescent tags will be removed, and hematoxylin and eosin (H&E) stains will be performed on the same tissue section. H&E slides were scanned at 20x magnification and co-registered with the fluorescent images using the Ultivue Ultistacker software.

#### Construction of the training data

We focused on the 34 non-lymph node tissue slides from the MSKCC cohort for training the model, 29 slides of which were used to generate the training image dataset. Depending on the number of positive cells for each marker, a fraction of the positive cells was randomly selected such that the training image dataset contained at least 50,000 positive cells. The same number of cells was selected randomly from the cells with negative labels from the same mIF slide to form the negative training image dataset. Therefore, the dataset for each marker consisted of over 100,000 positive and negative cells (50,000 each) from 29 whole mIF slides. The CellGate pipeline was trained on a dataset with over 750,000 single cells in total.

In the test dataset, regardless of the label for each cell, all cells in the 5 whole slide mIF images were used to test and evaluate the performance of the CellGate pipeline on the completely unseen whole slide mIF images. To diversify the whole slide mIF test samples, we divided the 38 mIF slides into 4 categories: 1. slides with a high number of cells (>75,000) and a high percentage of positive cells for the biomarker (>5%), 2. mIF slides with a relatively high number cells (15,000<cells<75,000) and a low percentage of positive cells (<5%), 3. mIF slides a low number of cells (<15,000) but a high percentage of positive cells (>5%), and 4. mIF slides with a low number of cells (<15,000) and a low percentage of positive cells (<5%). The unseen mIF test slides were selected randomly from these 4 categories. As a result, the unseen test slides were very diverse and the number of cells for each slide ranged between 7,000 to 403,000. A GPU cluster with 2 Nvidia A40 allowed us to train on hundreds of thousands of single cell images from whole mIF slides within hours.

#### Single cell image crop and processing

The cell image crops implemented to train the CNN models in CellGate included the cell borders and their immediate neighbors. The maximum and minimum height and width of the cell nuclei (X, Y) were obtained using the labeled dataset and were expanded for 6 pixels (≈ 2 µm) around the nucleus before reshaping the cell images to 64*64 size. Then a linear normalization for images was used in which all pixel values were divided by 65535.0 (for 16-bit images) before feeding them to the CNN models.

#### Gating the mIF images in the validation cohort (UNC samples)

The CellGate pipeline was first applied to the DAPI channel of the mIF and we performed the cell nuclei segmentation over the whole testing slides. In the next step, the cell coordinates were used to get the single cell images, and then the cell images were passed to the classification stage of the pipeline. The corresponding CNN models for each biomarker panel were applied to the cell images for that specific panel and then, we acquired the cell classification results for all cells across the mIF slides with the binary classification for each of the 8 biomarker panels. The output data from the CellGate pipeline also included the cell coordinates, as well as classification probabilities for each biomarker panel. Since this data is not gated, we used a previously developed data visualization method called “pseudo plots”.

## Supplemental Information

**Supplementary Figure 1.**
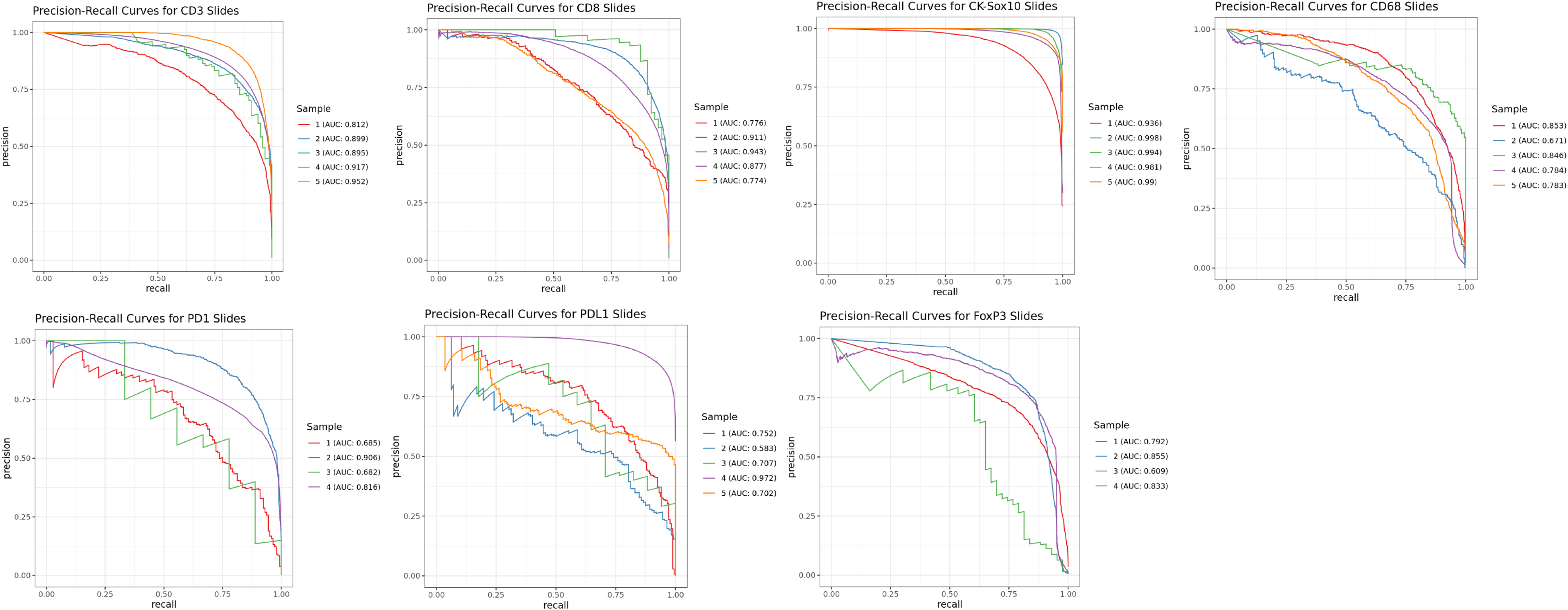
Precision-Recall Curves for unseen testing mIF whole slide for CD3, CD8, CK-Sox10, CD68, PD-1, PD-L1, and FOXP3 biomarkers.

**Supplementary Figure 2.**
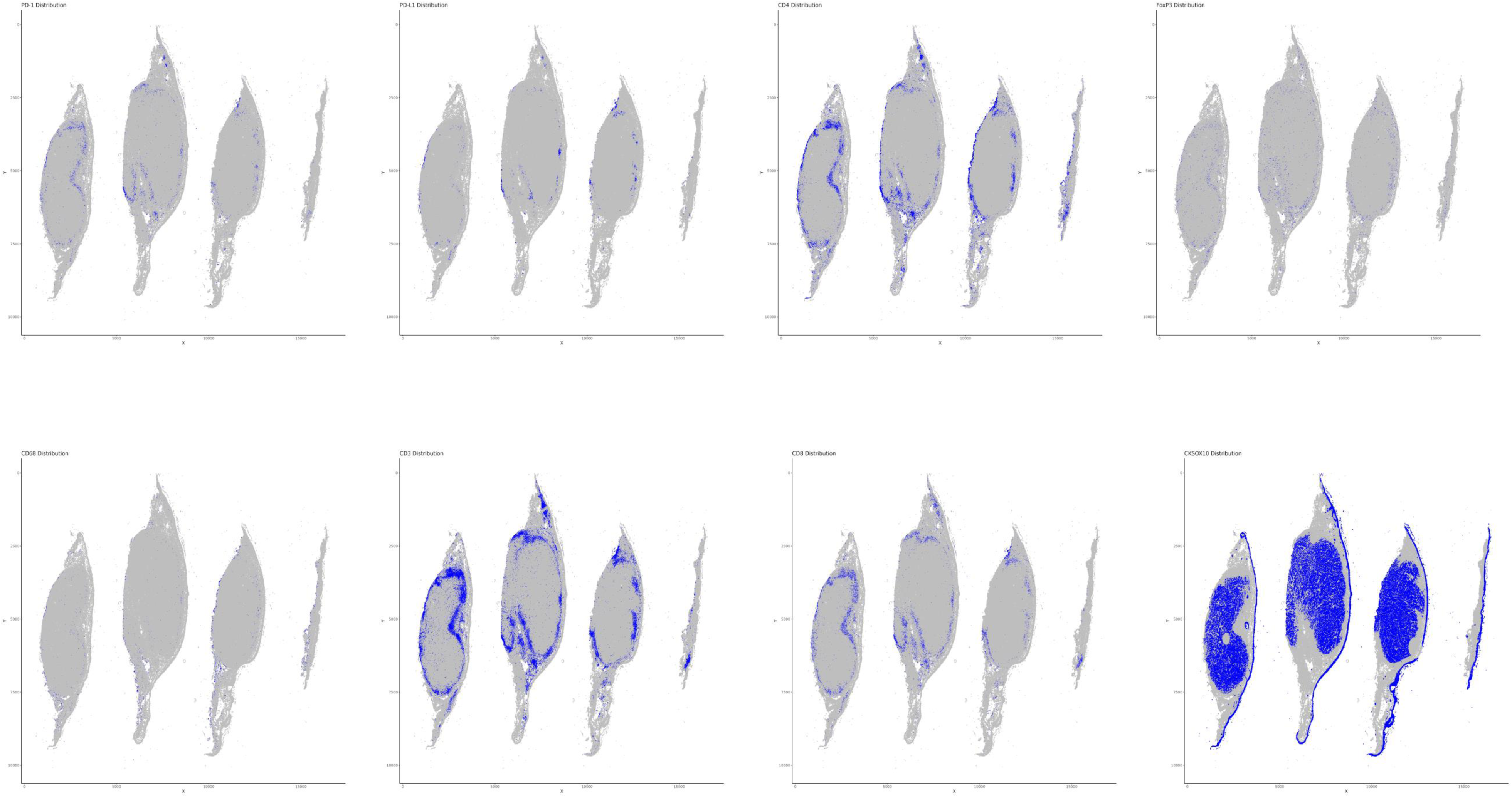
Binary classification of PD-1, PD-L1, CD4, FOXP3, CD68, CD3, CD8, and CK-Sox10 biomarker panels for the mIF whole slide image for UNC sample 1.

**Supplementary Figure 3.**
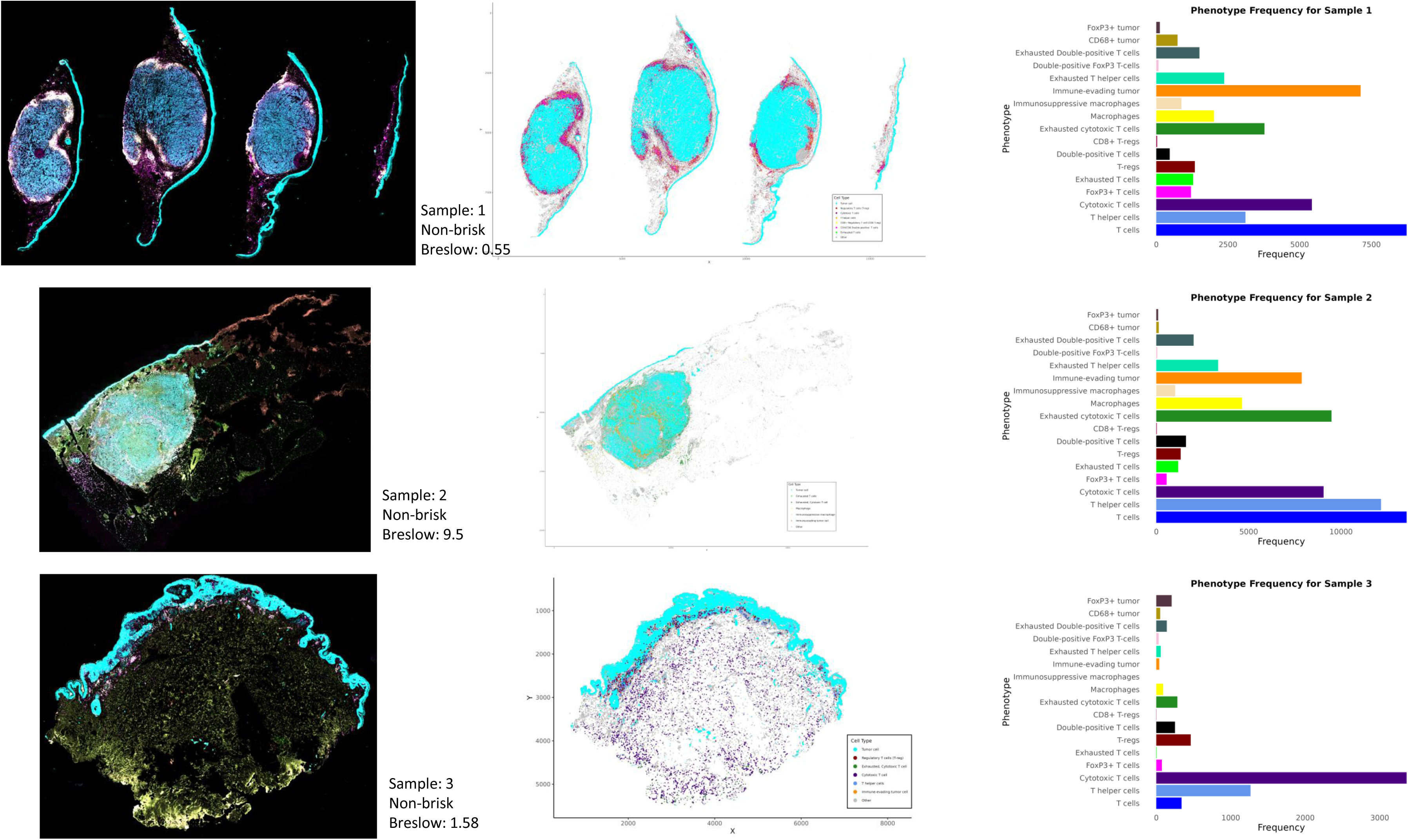
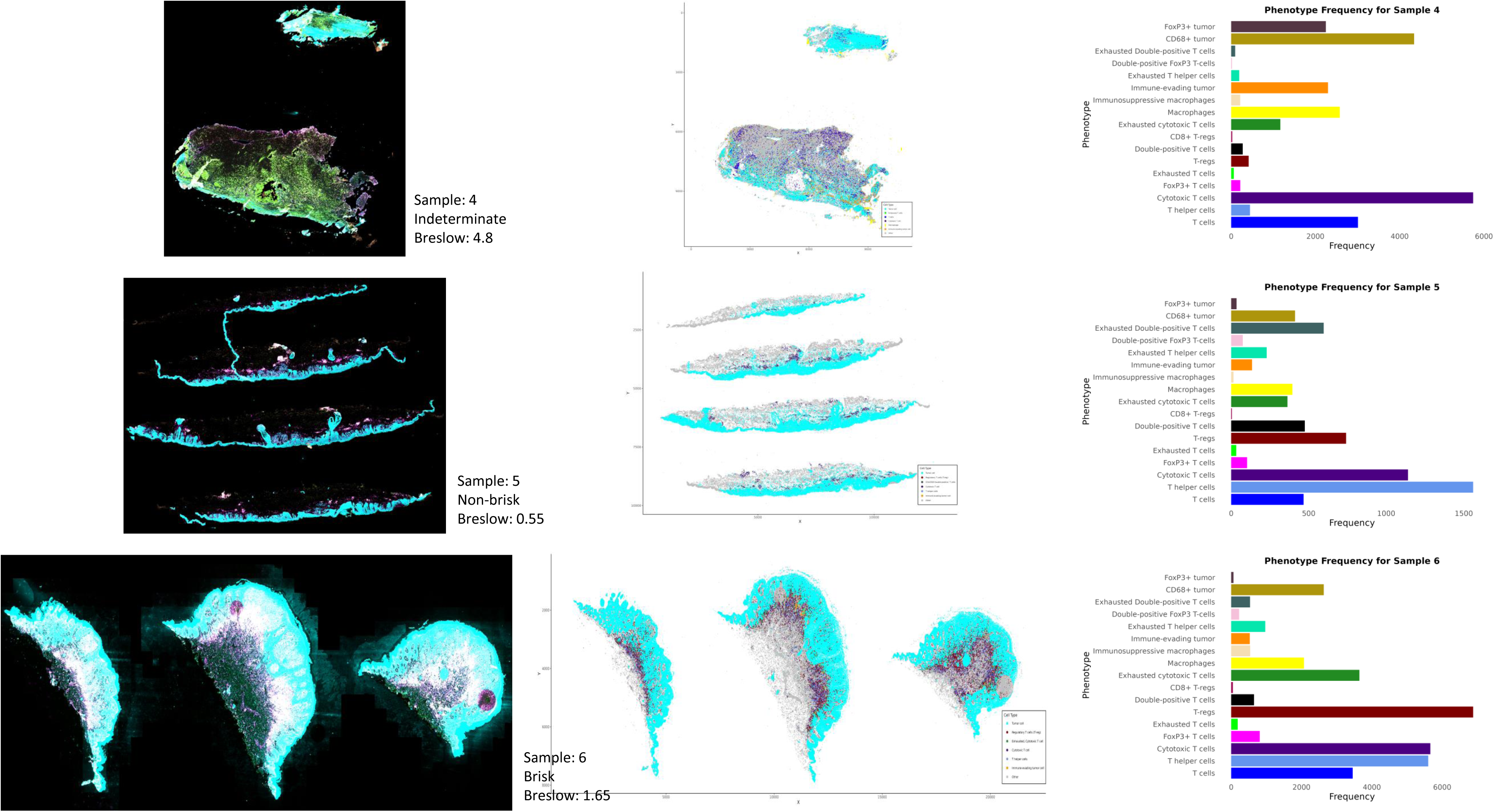
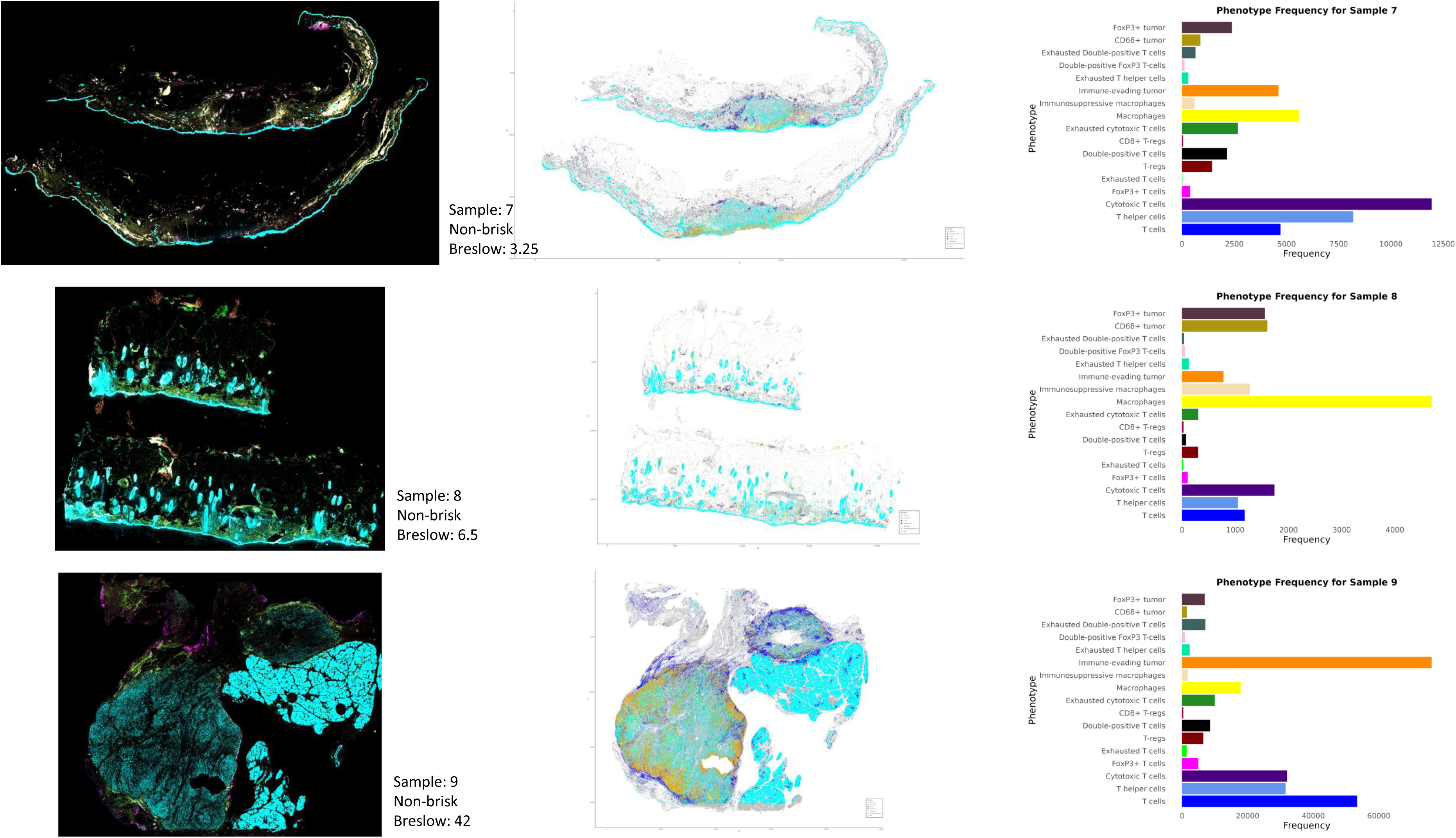
Pseudo plots and phenotype frequency bar charts for UNC mIF whole slide image samples 1-9.

**Supplementary Figure 4.**
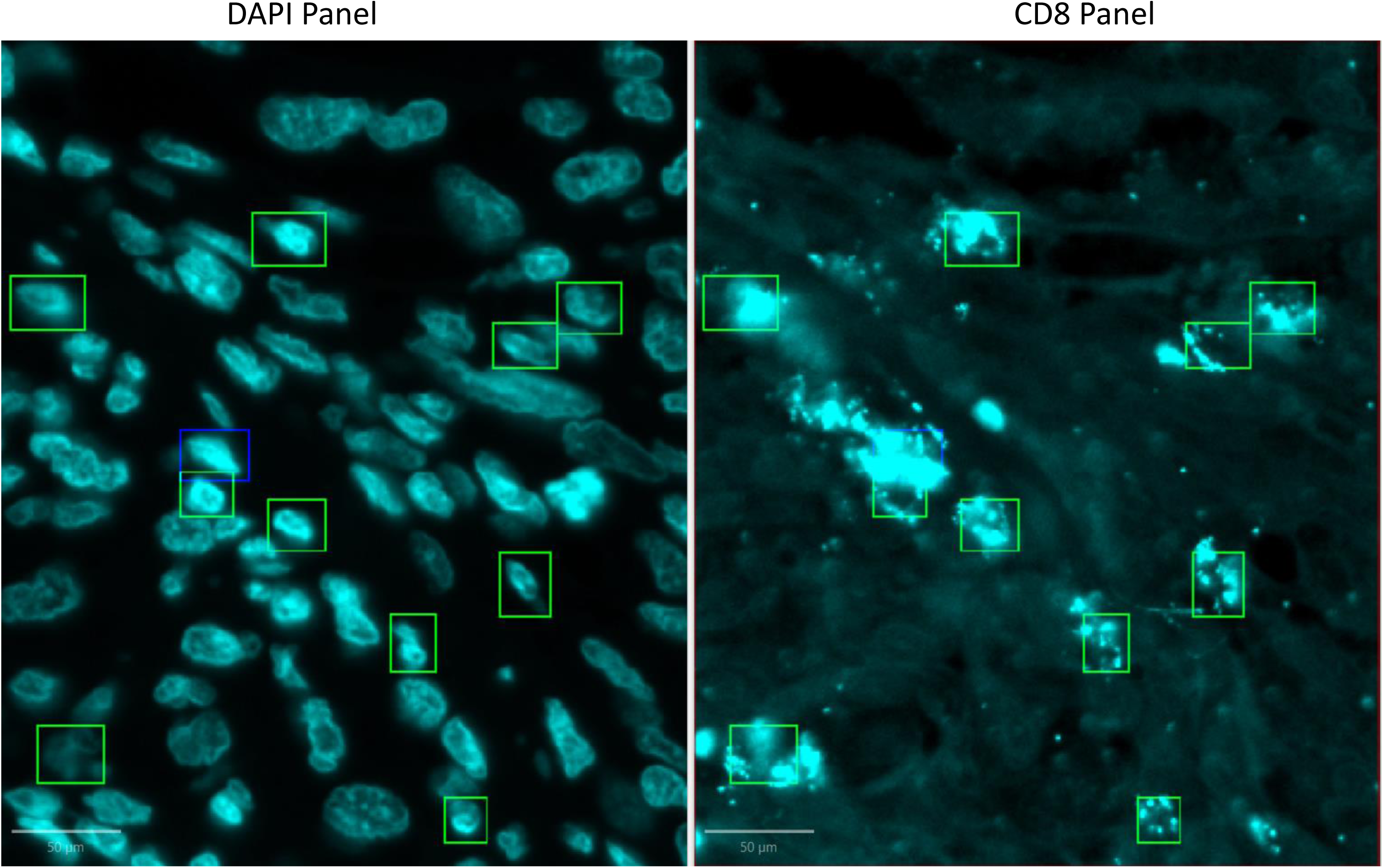
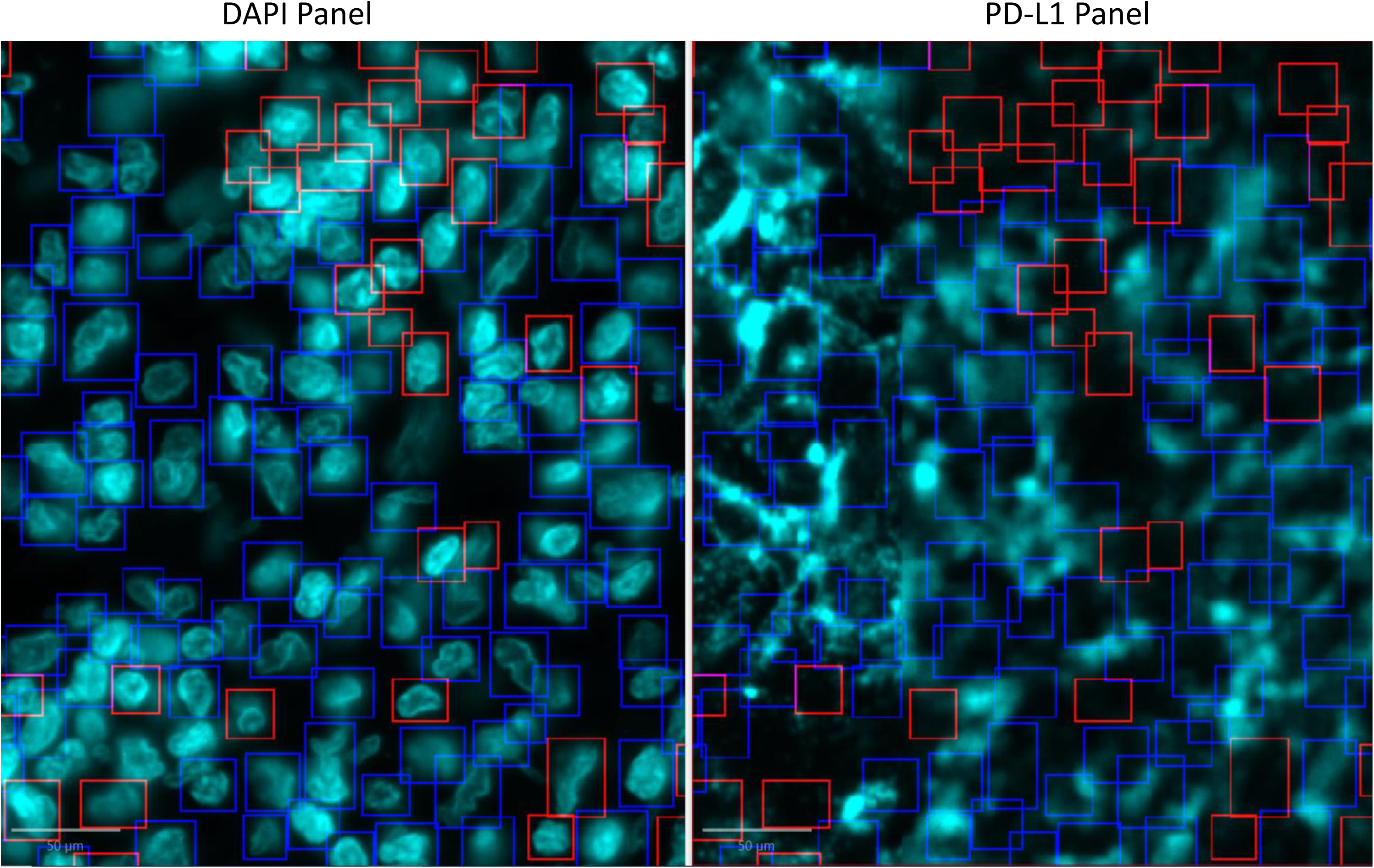
Region of interest of unseen testing slide 4 from raw mIF image. **A.** DAPI shows where the cells are located and the overlapping CD8 panel with the same coordinates is displayed. Green squares show the cells that were gated negative for CD8 biomarker originally, but gated positive by CellGate pipeline, showing the increase in sensitivity of the model due to considering the staining distribution within the whole cell. **B.** DAPI panel shows the nuclei and the PD-L1 displays the overlapping PD-L1 panel with the same coordinates. The blue squares show positive detection for both manual gating labels and the labels from the CellGate pipeline. Red squares are the cells that initially were classified positive using manual thresholding but labeled negative by the CellGate pipeline as the signals for these cells are at the background level intensity.

**Supplementary Table 1.** The CNN model performance metrics for CD3, CD8, CK-SOX10, CD68, PD-1, PD-L1, and FOXP3 biomarker panels on MSKCC unseen testing, whole slide mIF samples 1-5.

| A. CNN model performance metrics for CD3 Biomarker |  |  |  |  |  |  |  |  |
| --- | --- | --- | --- | --- | --- | --- | --- | --- |
| Sample | PPV | NPV | Sensitivity | Specificity | F-Score | Total Cells | CD3+ True Prop | CD3+ Pred Prop |
| 1 | 0.406 | 0.999 | 0.972 | 0.957 | 0.573 | 68788 | 0.029 | 0.070 |
| 2 | 0.476 | 1.000 | 0.993 | 0.987 | 0.643 | 315422 | 0.011 | 0.024 |
| 3 | 0.340 | 1.000 | 1.000 | 0.973 | 0.508 | 7160 | 0.014 | 0.041 |
| 4 | 0.905 | 0.859 | 0.717 | 0.958 | 0.801 | 403967 | 0.357 | 0.283 |
| 5 | 0.893 | 0.969 | 0.867 | 0.976 | 0.880 | 22919 | 0.190 | 0.184 |

| C. CNN model performance metrics for CK-Sox10 Biomarker |  |  |  |  |  |  |  |  |
| --- | --- | --- | --- | --- | --- | --- | --- | --- |
| Sample | PPV | NPV | Sensitivity | Specificity | F-Score | Total Cells | CK-Sox10+ True Prop | CK-Sox10+ Pred Prop |
| 1 | 0.624 | 0.991 | 0.978 | 0.809 | 0.762 | 68788 | 0.245 | 0.384 |
| 2 | 0.922 | 0.952 | 0.995 | 0.538 | 0.957 | 315422 | 0.845 | 0.912 |
| 3 | 0.995 | 0.763 | 0.887 | 0.988 | 0.938 | 7160 | 0.730 | 0.650 |
| 4 | 0.953 | 0.955 | 0.891 | 0.981 | 0.921 | 403967 | 0.301 | 0.281 |
| 5 | 0.980 | 0.854 | 0.868 | 0.978 | 0.921 | 22919 | 0.558 | 0.494 |

| E. CNN model performance metrics for PD-1 Biomarker |  |  |  |  |  |  |  |  |
| --- | --- | --- | --- | --- | --- | --- | --- | --- |
| Sample | PPV | NPV | Sensitivity | Specificity | F-Score | Total Cells | PD1+ True Prop | PD1+ Pred Prop |
| 1 | 0.287 | 1.000 | 0.924 | 0.995 | 0.438 | 68788 | 0.002 | 0.007 |
| 2 | 0.126 | 1.000 | 1.000 | 0.979 | 0.224 | 315422 | 0.003 | 0.024 |
| 3 | 0.114 | 1.000 | 1.000 | 0.990 | 0.205 | 7160 | 0.001 | 0.011 |
| 4 | 0.816 | 0.903 | 0.576 | 0.968 | 0.675 | 403967 | 0.197 | 0.139 |

| B. CNN model performance metrics for CD8 Biomarker |  |  |  |  |  |  |  |  |
| --- | --- | --- | --- | --- | --- | --- | --- | --- |
| Sample | PPV | NPV | Sensitivity | Specificity | F-Score | Total Cells | CD8+ True Prop | CD8+ Pred Prop |
| 1 | 0.309 | 1.000 | 0.992 | 0.980 | 0.472 | 68788 | 0.009 | 0.028 |
| 2 | 0.376 | 1.000 | 0.999 | 0.987 | 0.546 | 315422 | 0.008 | 0.021 |
| 3 | 0.253 | 1.000 | 1.000 | 0.973 | 0.404 | 7160 | 0.009 | 0.036 |
| 4 | 0.228 | 1.000 | 0.999 | 0.832 | 0.372 | 403967 | 0.047 | 0.208 |
| 5 | 0.276 | 0.998 | 0.979 | 0.795 | 0.431 | 22919 | 0.074 | 0.263 |

| D. CNN model performance metrics for CD68 Biomarker |  |  |  |  |  |  |  |  |
| --- | --- | --- | --- | --- | --- | --- | --- | --- |
| Sample | PPV | NPV | Sensitivity | Specificity | F-Score | Total Cells | CD68+ True Prop | CD68+ Pred Prop |
| 1 | 0.246 | 1.000 | 0.987 | 0.934 | 0.394 | 68788 | 0.021 | 0.086 |
| 2 | 0.215 | 1.000 | 0.958 | 0.997 | 0.351 | 315422 | 0.001 | 0.004 |
| 3 | 0.339 | 1.000 | 1.000 | 0.952 | 0.507 | 7160 | 0.024 | 0.070 |
| 4 | 0.647 | 0.998 | 0.823 | 0.994 | 0.724 | 403967 | 0.013 | 0.017 |
| 5 | 0.762 | 0.969 | 0.654 | 0.982 | 0.704 | 22919 | 0.082 | 0.071 |

| F. CNN model performance metrics for PD-L1 Biomarker |  |  |  |  |  |  |  |  |
| --- | --- | --- | --- | --- | --- | --- | --- | --- |
| Sample | PPV | NPV | Sensitivity | Specificity | F-Score | Total Cells | PDL1+ True Prop | PDL1+ Pred Prop |
| 1 | 0.085 | 1.000 | 0.989 | 0.973 | 0.157 | 68788 | 0.003 | 0.029 |
| 2 | 0.048 | 1.000 | 1.000 | 0.993 | 0.092 | 315422 | 0.000 | 0.007 |
| 3 | 0.053 | 1.000 | 1.000 | 0.958 | 0.101 | 7160 | 0.002 | 0.044 |
| 4 | 0.995 | 0.617 | 0.523 | 0.996 | 0.686 | 403967 | 0.565 | 0.298 |
| 5 | 0.150 | 1.000 | 1.000 | 0.958 | 0.261 | 22919 | 0.007 | 0.049 |

Supplementary Table 1. G. CNN model performance metrics for FOXP3 Biomarker
| Sample | PPV | NPV | Sensitivity | Specificity | F-Score | Total Cells | FoxP3+ True Prop | FoxP3+ Pred Prop |
| --- | --- | --- | --- | --- | --- | --- | --- | --- |
| 1 | 0.414 | 0.998 | 0.951 | 0.951 | 0.577 | 68788 | 0.035 | 0.080 |
| 2 | 0.298 | 1.000 | 0.944 | 0.988 | 0.453 | 315422 | 0.005 | 0.017 |
| 3 | 0.044 | 1.000 | 0.977 | 0.872 | 0.084 | 7160 | 0.006 | 0.133 |
| 4 | 0.520 | 1.000 | 0.948 | 0.994 | 0.672 | 403967 | 0.006 | 0.012 |

## Acknowledgment

This work is supported in part by the MSKCC Society, the V Foundation, the Parker Institute for Cancer Immunotherapy, NIH P30 CA008748, NIH R01 CA276286, NIH P01 CA206980, NIH R01 CA233524, and the MSK-MIND consortium.

## Notes

### Competing Interest Statement

The authors have declared no competing interest.

## Reference

1. Giesen, C. et al. Highly multiplexed imaging of tumor tissues with subcellular resolution by mass cytometry. Nat Methods 11, 417–22 (2014).

2. Goltsev, Y. et al. Deep Profiling of Mouse Splenic Architecture with CODEX Multiplexed Imaging. Cell 174, 968–981 e15 (2018).

3. Berry, S. et al. Analysis of multispectral imaging with the AstroPath platform informs efficacy of PD-1 blockade. Science 372(2021).

4. Adegoke, N.A. et al. Classification of the tumor immune microenvironment and associations with outcomes in patients with metastatic melanoma treated with immunotherapies. J Immunother Cancer 11(2023).

5. Lin, J.R. et al. Highly multiplexed immunofluorescence imaging of human tissues and tumors using t-CyCIF and conventional optical microscopes. Elife 7(2018).

6. Gut, G., Herrmann, M.D. & Pelkmans, L. Multiplexed protein maps link subcellular organization to cellular states. Science 361(2018).

7. Tsujikawa, T. et al. Quantitative Multiplex Immunohistochemistry Reveals Myeloid-Inflamed Tumor-Immune Complexity Associated with Poor Prognosis. Cell Rep 19, 203–217 (2017).

8. Ptacek, J. et al. Multiplexed ion beam imaging (MIBI) for characterization of the tumor microenvironment across tumor types. Lab Invest 100, 1111–1123 (2020).

9. Lu, S. et al. Comparison of Biomarker Modalities for Predicting Response to PD-1/PD-L1 Checkpoint Blockade: A Systematic Review and Meta-analysis. JAMA Oncol 5, 1195–1204 (2019).

10. Lin, J.R. et al. High-plex immunofluorescence imaging and traditional histology of the same tissue section for discovering image-based biomarkers. Nat Cancer 4, 1036–1052 (2023).

11. Greenwald, N.F. et al. Whole-cell segmentation of tissue images with human-level performance using large-scale data annotation and deep learning. Nature biotechnology 40, 555–565 (2022).

12. Stringer, C., Wang, T., Michaelos, M. & Pachitariu, M. Cellpose: a generalist algorithm for cellular segmentation. Nature methods 18, 100–106 (2021).

13. Lee, M.Y. et al. CellSeg: a robust, pre-trained nucleus segmentation and pixel quantification software for highly multiplexed fluorescence images. BMC bioinformatics 23, 46 (2022).

14. Schmidt, U., Weigert, M., Broaddus, C. & Myers, G. Cell detection with star-convex polygons. in Medical Image Computing and Computer Assisted Intervention–MICCAI 2018: 21st International Conference, Granada, Spain, September 16-20, 2018, Proceedings, Part II 11 265–273 (Springer, 2018).

15. Yapp, C. et al. UnMICST: Deep learning with real augmentation for robust segmentation of highly multiplexed images of human tissues. Communications Biology 5, 1263 (2022).

16. Van Gassen, S. et al. FlowSOM: Using self-organizing maps for visualization and interpretation of cytometry data. Cytometry Part A 87, 636–645 (2015).

17. Liu, C.C. et al. Robust phenotyping of highly multiplexed tissue imaging data using pixel-level clustering. Nature Communications 14, 4618 (2023).

18. Geuenich, M.J., Hou, J., Lee, S., Ayub, S., Jackson, H.W. & Campbell, K.R. Automated assignment of cell identity from single-cell multiplexed imaging and proteomic data. Cell Systems 12, 1173–1186. e5 (2021).

19. Brbić, M. et al. Annotation of spatially resolved single-cell data with STELLAR. Nature Methods 19, 1411–1418 (2022).

20. Yosofvand, M. et al. Automated detection and scoring of tumor-infiltrating lymphocytes in breast cancer histopathology slides. Cancers 15, 3635 (2023).

21. Amitay, Y., Bussi, Y., Feinstein, B., Bagon, S., Milo, I. & Keren, L. CellSighter: a neural network to classify cells in highly multiplexed images. Nature communications 14, 4302 (2023).

22. Bhinder, B., Gilvary, C., Madhukar, N.S. & Elemento, O. Artificial Intelligence in Cancer Research and Precision Medicine. Cancer Discov 11, 900–915 (2021).

23. Ehteshami Bejnordi, B., et al. Diagnostic Assessment of Deep Learning Algorithms for Detection of Lymph Node Metastases in Women With Breast Cancer. JAMA 318, 2199–2210 (2017).

24. Coudray, N. et al. Classification and mutation prediction from non-small cell lung cancer histopathology images using deep learning. Nat Med 24, 1559–1567 (2018).

25. Simonyan, K. & Zisserman, A. Very deep convolutional networks for large-scale image recognition. arXiv preprint arXiv: 1409.1556 (2014).

26. Deng, J., Dong, W., Socher, R., Li, L.-J., Li, K. & Fei-Fei, L. Imagenet: A large-scale hierarchical image database. in 2009 IEEE conference on computer vision and pattern recognition 248–255 (Ieee, 2009).

27. Abadi, M. et al. {TensorFlow}: a system for {Large-Scale} machine learning. in 12th USENIX symposium on operating systems design and implementation (OSDI 16) 265–283 (2016).

28. Smithy, J.W. et al. Spatial assessment of stromal B cell aggregates predicts response to checkpoint inhibitors in unresectable melanoma. medRxiv, 2024.08.09.24311758 (2024).

29. Vanguri, R.S. et al. Integration of peripheral blood– and tissue-based biomarkers of response to immune checkpoint blockade in urothelial carcinoma. J Pathol 261, 349–360 (2023).

30. Thomas, N.E. et al. Tumor-infiltrating lymphocyte grade in primary melanomas is independently associated with melanoma-specific survival in the population-based genes, environment and melanoma study. J Clin Oncol 31, 4252–9 (2013).

31. Tumeh, P.C. et al. PD-1 blockade induces responses by inhibiting adaptive immune resistance. Nature 515, 568–71 (2014).

32. Taube, J.M. et al. Association of PD-1, PD-1 ligands, and other features of the tumor immune microenvironment with response to anti-PD-1 therapy. Clin Cancer Res 20, 5064–74 (2014).

33. Attrill, G.H. et al. Detailed spatial immunophenotyping of primary melanomas reveals immune cell subpopulations associated with patient outcome. Front Immunol 13, 979993 (2022).

34. Schad, S.E. et al. Tumor-induced double positive T cells display distinct lineage commitment mechanisms and functions. J Exp Med 219(2022).

35. Xie, F., Xu, M., Lu, J., Mao, L. & Wang, S. The role of exosomal PD-L1 in tumor progression and immunotherapy. Mol Cancer 18, 146 (2019).

36. Chen, J. et al. Tumor extracellular vesicles mediate anti-PD-L1 therapy resistance by decoying anti-PD-L1. Cell Mol Immunol 19, 1290–1301 (2022).

37. Chen, G. et al. Exosomal PD-L1 contributes to immunosuppression and is associated with anti-PD-1 response. Nature 560, 382–386 (2018).

38. Poggio, M. et al. Suppression of Exosomal PD-L1 Induces Systemic Anti-tumor Immunity and Memory. Cell 177, 414–427 e13 (2019).

39. Kuett, L. et al. Three-dimensional imaging mass cytometry for highly multiplexed molecular and cellular mapping of tissues and the tumor microenvironment. Nat Cancer 3, 122–133 (2022).

40. Lin, J.R. et al. Multiplexed 3D atlas of state transitions and immune interaction in colorectal cancer. Cell 186, 363–381 e19 (2023).

41. Antoranz, A. et al. Mapping the Immune Landscape in Metastatic Melanoma Reveals Localized Cell-Cell Interactions That Predict Immunotherapy Response. Cancer Res 82, 3275–3290 (2022).

42. Phillips, D. et al. Immune cell topography predicts response to PD-1 blockade in cutaneous T cell lymphoma. Nat Commun 12, 6726 (2021).

43. Vanhersecke, L. et al. Mature tertiary lymphoid structures predict immune checkpoint inhibitor efficacy in solid tumors independently of PD-L1 expression. Nat Cancer 2, 794–802 (2021).

44. Italiano, A. et al. Pembrolizumab in soft-tissue sarcomas with tertiary lymphoid structures: a phase 2 PEMBROSARC trial cohort. Nat Med 28, 1199–1206 (2022).

45. Brunet, M. et al. Mature tertiary lymphoid structure is a specific biomarker of cancer immunotherapy and does not predict outcome to chemotherapy in non-small-cell lung cancer. Ann Oncol 33, 1084–1085 (2022).

